# Rv3400 is a phosphoglucomutase required for trehalose metabolism in *Mycobacterium tuberculosis*

**DOI:** 10.1101/2025.11.20.689507

**Authors:** Yi Liu, Nadine Ruecker, Valwynne Faulkner, Kavitha Rachineni, Nada Al-Saffar, Pablo Villacampa Teixeira, Tiago R D Costa, Eachan Johnson, Brian D. Robertson, Sabine Ehrt, Gerald Larrouy-Maumus

## Abstract

*Mycobacterium tuberculosis* is a global killer causing over a million deaths from tuberculosis (TB) every year. It is therefore a major burden on human health. To reduce the deadly impact of TB, we need a better understanding of the strategies used by *M. tuberculosis* to adapt its metabolism to survive and persist in the human host. This will help us design better strategies for TB treatment and control. Previous enzymological studies have reported the Mtb *rv3400* gene as encoding a β-phosphoglucomutase; however, its role in *M. tuberculosis* metabolism was not investigated. In the present study, we show that deletion of *rv3400* leads to a 30-fold increase in β-D-glucose 1-phosphate, confirming its primary function as a β-phosphoglucomutase. Additionally, deletion of *rv3400* leads to a growth defect when trehalose is the sole carbon source indicting that this enzyme is required for optimal use of trehalose, an essential disaccharide in mycobacteria.

**Importance:** Trehalose metabolism plays a corner stone in Mycobacterium tuberculosis physiology and virulence. Hence, a better understanding of the metabolism of this essential disaccharide is required to develop novel strategies to eradicate Tuberculosis. Here we report on the characterisation of the strain lacking a β-phosphoglucomutase, encoded by the gene *rv3400*. We show that deletion of *rv3400* leads to an increase of β-D-glucose 1-phosphate and a growth defect when using trehalose as sole carbon source. Taken together, the data presented here provide evidence that Rv3400 is required for the catabolism of trehalose.

## Introduction

*Mycobacterium tuberculosis* (Mtb) causes over 1 million deaths annually, maintaining its status as a major global health threat (1). Although antibiotics have been used for decades, drug resistance and persistent infections severely limit current treatments against this global killer. A key challenge in combating Mtb lies in addressing its ability to adapt to host environments, such as macrophages, by reprogramming its metabolism and gene expression. A comprehensive understanding of this knowledge gap is important for the identification of novel druggable targets and for achieving the WHO’s tuberculosis eradication goals through next-generation therapeutics.

Recent *in vitro* studies have characterised Rv3400 as a β-phosphoglucomutase (β-PGM), an enzyme that catalyses the isomerization of β-D-glucose-1-phosphate (G1P) to β -D-glucose-6-phosphate (G6P) via a β-D-glucose-1,6-bisphosphate (β-G1,6bisP) intermediate (2). The function of this enzyme is closely linked to central carbon metabolism, as G6P serves as a key metabolic node for the production of ATP and NADPH via glycolysis and the pentose phosphate pathway, respectively (3). Although dispensable for *in vitro* growth of Mtb H37Rv (4-6), Rv3400 is required for bacterial growth in the spleen of C57BL/6J mice (5, 6). Furthermore, using a genetic interaction mapping strategy, Joshi et al. reported the interaction of Rv3400 with the virulence-associated mycobacterial cell entry 4 (*mce4*) operon, indicating its potential involvement in Mtb virulence (7).

Recent studies also suggest that Rv3400 could be part of the trehalose degradation pathway(2). Trehalose is a non-reducing disaccharide (1-O-α-D-glucopyranosyl-α-D-glucopyranoside) found in insects, plants and microbial cells. In mycobacteria, trehalose is constitutively present in cells grown in standard laboratory media and is the only free sugar readily detectable in the cytoplasm, comprising approximately 1.5–3% of the total cellular dry weight (8). Trehalose has well-established protective functions during cryopreservation and desiccation and is known to enhance microbial survival under environmental stress *in vivo*. In mycobacteria, trehalose is not only a reserved energy source, but also an active player in metabolic pathways, as suggested by the dynamic turnover of the trehalose pool found in the rapid-growing mycobacteria *Mycobacterium smegmatis* (9). It is incorporated into many mycobacterial glycolipids, including cord factor (trehalose 6,6’-dimycolate), sulpholipids (acylated trehalose-2’-sulphate derivatives) and trehalose-containing lipooligosaccharides (10-12). The multiple roles of these metabolites further underscore the importance of trehalose metabolism in Mtb survival and pathogenesis.

Here, we investigated the impact of *rv3400* deletion on Mtb metabolism and found that the Mtb Δ*rv3400* strain has a growth defect on trehalose with the accumulation of β-D-glucose-1-phosphate, consistent with the loss of β-PGM activity.

## Results and Discussion

### Deletion of *rv3400* leads to an increase in β-D-glucose 1-phosphate level

Recent work has identified Rv3400 as a β-PGM *in vitro*(2). To confirm its enzymatic function, we constructed the *rv3400* deletion mutant (Δ*rv3400*), complemented (Δ*rv3400*::*rv3400*) and catalytic site mutant (Δ*rv3400*::*rv3400*_D29A_) strains and measured their growth in carbon-defined liquid medium. As shown in Figure 1A, compared to the parental strain, deletion of *rv3400* did not impair growth when glucose was used as sole carbon source. To further validate the role of Rv3400 as a β-PGM, targeted metabolomics analysis revealed five peaks in the extracted ion chromatogram for ion at *m*/*z* 259.024 [M-H]^-^, assigned to monosaccharide phosphates: α-D-fructose 6-phophate (retention time (RT) 11.1 minutes), α-D-glucose-1-phosphate (RT 11.3 minutes), α-D-mannose-1-phosphate/ α-D-mannose-6-phosphate (RT 11.6 minutes), β-D-glucose-1-phosphate (RT 11.8 minutes) and α-D-glucose-6-phosphate (RT 12.4 minutes). In the Δ*rv3400* strain, the peak corresponding to β-D-glucose-1-phosphate (RT 11.8 minutes) increased (Figure 1B). The attribution of that peak assigned to β-D-glucose-1-phosphate was also confirmed by nuclear magnetic resonance (NMR) (Supporting text and figures 1-6). In addition, quantification of the 4 monosaccharide phosphates informed that *rv3400* deletion specifically elevated the concentration of β-D-glucose-1-phosphate (Figure 1C), confirming that Rv3400 is a β-PGM involved in the conversion of β-D-glucose-1-phosphate into α-D-glucose-6-phosphate.

**Figure 1:**
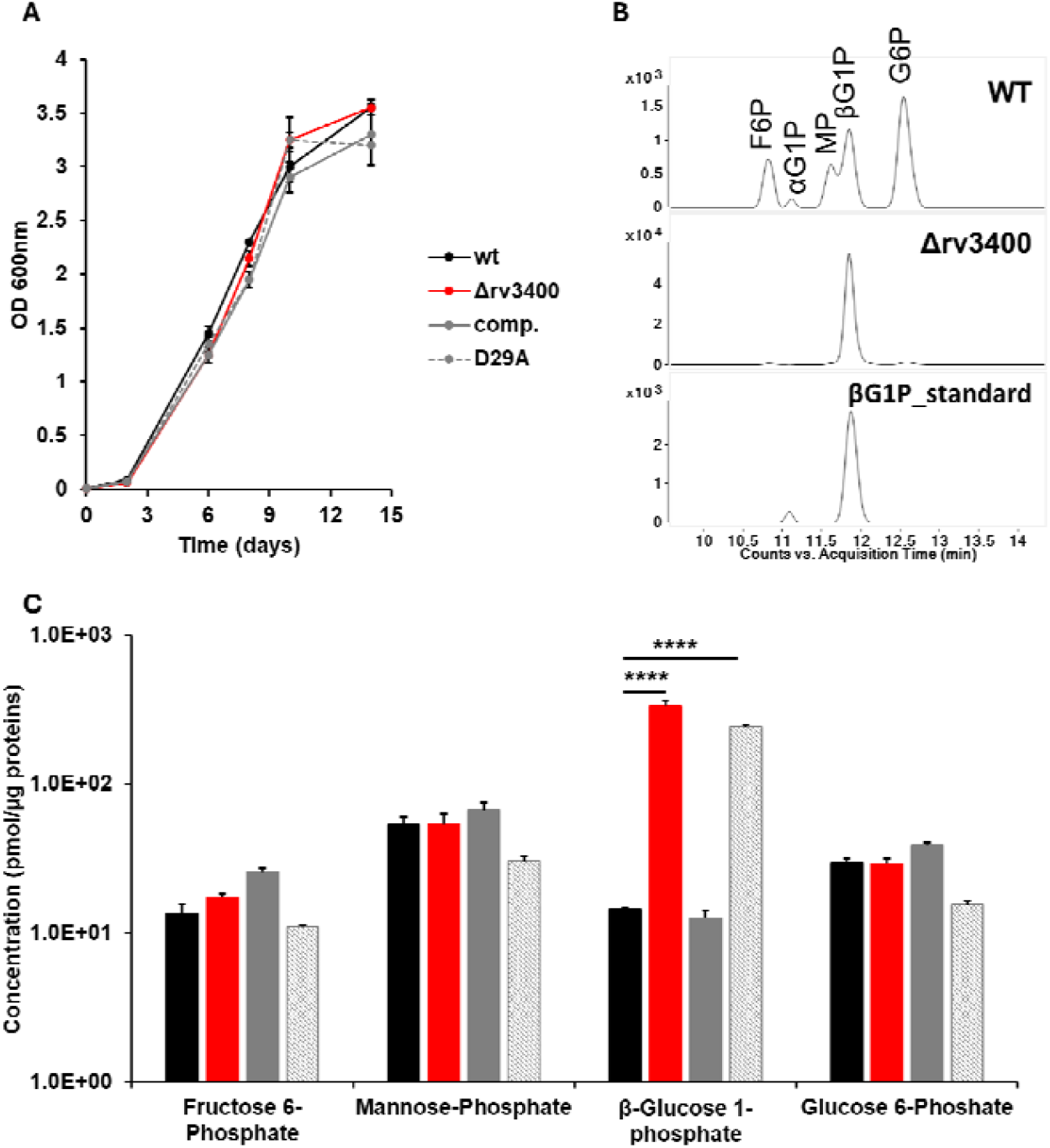
Rv3400 is a phosphoglucomutase in *Mycobacterium tuberculosis* H37Rv. (**A**). Growth curves of parental (black) and Δ*rv3400* (red) strains with glucose as the sole carbon source. (**B**). Extraction ion chromatograms for ion at *m*/*z* 259.024 [M-H]-for the parental (top), Δ*rv3400* strains (middle) and β-D-glucose-1-phosphate standard (bottom). (**C**). Bar charts displaying the concentration in pmol/µg proteins for the 5 relevant phosphorylated sugars (F6P: α-D-fructose-6-phosphate; αG1P: α-D-glucose-1-phosphate; βG1P: α-D-glucose-1-phosphate; MP: α-D-mannose-6-phosphate/ α-D-mannose-1-phosphate; G6P: α-D-glucose-6-phosphate). wild type: back bars, Δ*rv3400*: red bars, complemented: grey bars and D29A: shaded grey bars. Asterisks indicate levels of significance compared to parental strain bacteria; the significance level of differences were determined by unpaired Student’s t-tests, with *: *p* < 0.05; **: *p* < 0.01; ***: *p* < 0.001; ****: *p* < 0.0001.

### Deletion of *rv3400* leads to impaired growth when trehalose was used as the sole carbon source

The trehalose recycling pathway mediated by the LpqY-SugA-SugB-SugC ATP-binding cassette transporter is critical during acute Mtb infection in the mouse model when exponential replication occurs, as trehalose can serve as a carbon source; disruption of this pathway may cause starvation in nutrient-limited *in vivo* microenvironments (13, 14). Given Rv3400’s role in trehalose recycling, we assessed the growth patterns of Δ*rv3400* and Δ*rv3400*::*rv3400*_D29A_ strains with trehalose as the only carbon source. As shown in Figure 2A, both strains exhibited growth defects under this condition.

**Figure 2:**
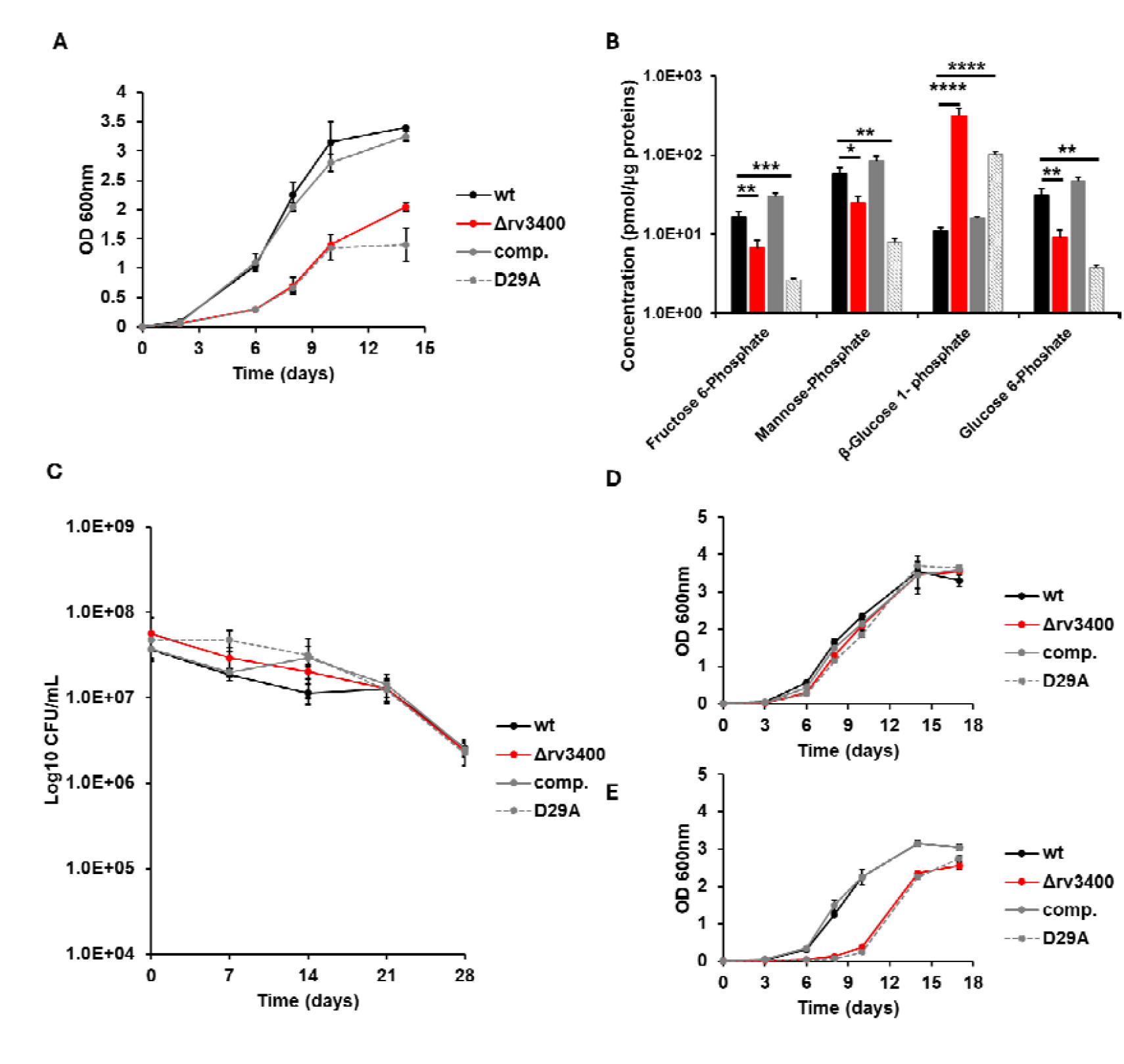
Rv3400 is required for trehalose recycling but dispensable during starvation. (**A**). Growth curves of parental (black), Δ*rv3400* (red), complemented (grey) and D29A (dashed grey) strains with trehalose as the sole carbon source. (**B**). Bar charts displaying the concentration in pmol/µg proteins for the 4 relevant phosphorylated sugars (F6P: α-D-fructose-6-phosphate; βG1P: α-D-glucose-1-phosphate; MP: α-D-mannose-6-phosphate/ α-D-mannose-1-phosphate; G6P: α-D-glucose-6-phosphate). wild type: back bars, Δ*rv3400*: red bars, completed: grey bars and D29A: shaded grey bars. (**C**). Survival assay under PBS starvation as measured by the number of viable cells (CFU/ml). (**D**) Regrowth in 7H9 glucose of parental (black), Δ*rv3400* (red), complemented (grey) and D29A (dashed grey) strains following 28 days starvation on PBS. (**E**) Regrowth in 7H9 trehalose of parental (black), Δ*rv3400* (red), complemented (grey) and D29A (dashed grey) strains following 28 days starvation on PBS. Asterisks indicate levels of significance compared to parental strain bacteria; the significance level of differences were determined by unpaired Student’s *t*-tests, with *: *p* < 0.05; **: *p* < 0.01; ***: *p* < 0.001; ****: *p* < 0.0001.

To further consolidate this observation, we conducted growth curves using 0.05%, 0.02% and 0.4% glucose or trehalose. As expected, we do observe an increase in growth when using higher concentration of carbon source in the wild-type strain and complemented strains, although no increase in growth was noticed when more carbon where available to the culture medium to the Δ*rv3400* strain (supplementary figure 6). In addition, to rule out any contaminant that could interfere with bacterial growth such as fatty acids originated from the standard BSA, we conducted the same experiment with BSA fraction V fatty acids free. A similar result is observed where higher concentrations of carbon source where the Δ*rv3400* strain does not display any increase in growth with increase trehalose concentration (supplementary figure 7). Taken together these data clearly demonstrate that Rv3400 is required for optimal use of trehalose in Mtb.

Targeted metabolomics further revealed that the Δ*rv3400* mutant accumulated a 30-fold higher β-D-glucose-1-phosphate pool, while showing 2-fold, 2-fold, and 3-fold reductions in α-D-fructose 6-phosphate, α-D-mannose 1-phosphate/6-phosphate, and α-D-glucose 6-phosphate pools, respectively (Figure 2B). A similar trend is observed in the metabolic profile of the D29A strain (Figure 2B). Further long-term PBS starvation assay showed that, although required for optimal growth when trehalose was the sole carbon source, Rv3400 is not required for bacterial viability during starvation (Figure 2C), but necessary for optimal recovery after long-term starvation when using trehalose as carbon source but not glucose (Figure 2D and Figure 2E). Taken together, these data indicate that Rv3400 is part of the trehalose recycling pathway and essential to sustain Mtb optimal growth when trehalose was used as sole carbon source.

Taken together, our data demonstrate that Rv3400 is a β-PGM required for optimal utilisation of trehalose as sole carbon sources. This study provides the first biochemical validation of Rv3400’s functions, highlighting its roles in Mtb resuscitation when trehalose serves as an important carbon source. The trehalose synthesis and recycling pathway has been studied for its involvement not only in Mtb regrowth but also in the formation of persisters and the development of multidrug resistance (15, 16). Therefore, our study on Rv3400 contributes substantially to understanding this pathway as an important part of Mtb metabolism and a compelling therapeutic target.

## Material and Methods

### Bacterial strains and growth conditions

The Mtb Δ*rv3400* mutant was constructed in strain H37Rv by allelic exchange using a recombineering approach as previously described (17).

A pMV306 plasmid with the *rv3400* native promoter was used as the backbone for construction of the *rv3400* complementation plasmid. To generate a complementation of *rv3400* under its native promoter, the integrating expression vector pMV306 KanR (18) was digested with EcoRV-HF (NEB) and subsequently column purified using the QIAquick purification kit (Qiagen). DNA fragments comprising the full length of *rv3400* and *rv3400* D29A along with 150 bp upstream of the annotated start codon were synthesised (GenScript). The synthesised fragments included 20 bp overhangs at both the 5’ and 3’ ends homologous to the digested vector. NEBuilder HiFi DNA assembly (NEB) was used to ligate the linearised vector and synthesised inserts. Correct assembly of the recombinant plasmids were confirmed by Nanopore sequencing.

Precultures of all strains were grown in 10 mL square bottles with 7H9 medium (Sigma) supplemented with 0.2% w/v glucose, 0.5% w/v standard bovine serum albumin (BSA), 0.08% w/v NaCl, 0.05% w/v tyloxapol and appropriate antibiotics. The bottles were incubated in a shaking incubator at 37 °C, 120 rpm. 7H10 agar plates (Sigma) were supplemented with 0.2% w/v glycerol, 0.2% w/v dextrose, 0.5% w/v BSA and 0.08% w/v NaCl for growth on solid medium in a 37°C static incubator. Fo carbon source concentration dependence, BSA fraction V fatty acid free was used, while carbon source concentrations were either 0, 0.05%, 0.2% or 0.4% final concentration w/v.

### Growth curves

Growth curve assays were performed in 10 mL square bottle (Nalgene) with pre-warmed 7H9 medium. Pre-cultures of bacteria were grown and used to inoculate these flasks at OD_600nm_ = 0.01. Inoculated flasks were incubated in a 37°C static incubator and OD_600nm_ measurements were taken at least 5 times to produce growth curves until the stationary phase.

### Bacteria viability under starvation

To quantify the number of viable cells under PBS starvation, mid-log phase cultures were spun down, washed twice with PBS and resuspended at a OD_600nm_ = 0.1 in PBS+0.5% Tyloxapol. Cultures were sampled at day 0, 7, 14, 21 and 28 and plated on 7H10 agar plates. Colonies were counted after 4 weeks incubation at 37 °C.

### Metabolite extraction

Mtb strains were grown in 10 mL flasks with 7H9 medium to reach OD_600nm_ of approximately 0.8. After incubation, bacteria were collected by centrifugation at 3,000 x g, 4 °C for 10 minutes. Pellets were washed once with cold sterile PBS and metabolically quenched in extraction solution (40% acetonitrile, 40% methanol, 20% ddH_2_O). Samples were transferred to microtubes containing 0.1 mm acid-washed zirconia beads and lysed using a FastPrep-24 homogeniser twice at 6.0 m/s for 30 seconds, with a 5-minute interval. Supernatants were transferred to 0.22 μm spin X column filters (Costar) and filtered twice by centrifuging at 15,000 x g for 30 minutes. After filtration, the flowthrough was mixed with equal volume of acetonitrile and centrifuged at 15,000 x g for 10 minutes. 100 µL of mixture was loaded into polypropylene snap vials (Agilent Technologies) for LC-MS analysis.

### Liquid chromatography-mass spectrometry (LC-MS)

The data were acquired with an Agilent 1290 Infinity II UHPLC coupled to a 6545 LC/Q-TOF system. Chromatographic separation was performed with an Agilent InfinityLab Poroshell 120 HILIC-Z (2.1 × 100 mm, 2.7 μm (p/n 675775-924)) column. The HILIC-Z methodology was optimized for polar acidic metabolites. Column compartment was set at 50°C. For easy and consistent mobile-phase preparation, a concentrated 10 x solution consisting of 100 mM ammonium acetate (pH 9.0) in water was prepared to produce mobile phases A and B. Mobile phase A consisted of 10 mM ammonium acetate in water (pH 9) with a 5 μM Agilent InfinityLab deactivator additive (p/n 5191-4506), and mobile phase B consisted of 10 mM ammonium acetate (pH 9) in 10:90 (v:v) water/acetonitrile with a 5 μM Agilent InfinityLab deactivator additive (p/n 5191-4506). The following gradient was applied at a flow rate of 0.25 ml/min: 0 min, 96% B; 2 min, 96% B; 5.5 min, 88% B; 8.5 min, 88% B; 9 min, 86% B; 14 min, 86% B; 17 min, 82% B; 23 min, 65% B; 24 min, 65% B; 24.5 min, 96% B; 26 min, 96% B and 3-min of re-equilibration at 96% B. Accurate MS was performed using an Agilent Accurate Mass 6545 QTOF apparatus. Dynamic mass axis calibration was achieved by continuous infusion after the chromatography of a reference mass solution using an isocratic pump connected to an electrospray ionization source operated in negative-ion mode. The following parameters were used: gas temperature, 225°C; drying gas, 13 l min^-1^; sheath gas temperature, 350 °C; nebulizer pressure, 35 psi; sheath gas flow, 12 l min^-1^; capillary voltage, 3,500 V; nozzle voltage, 0 V; fragmentor voltage, 125 V; skimmer 45V and octupole 1 RF voltage, 750V. The data were collected in centroid 4 GHz (extended dynamic range) mode.

### LC-MS metabolomics data analysis

MassHunter Qualitative Analysis software was used to check the extracted ion chromatograms of the *m/z* values of interest and obtain their retention times, which were used to construct the personal compound database and library (PCDL) for following analysis. MassHunter Profinder B8.0 was used to extract the stable isotope labelling patterns from PCDL. Standards were purchased from Sigma-Aldrich: α-D-fructose-6-phosphate (F1502-1G), α-D-glucose-6-phosphate (G7250), α-D-glucose-1-phosphate (G7000-5G), calbiochem α-D-mannose-6-phosphate (444100-50MG) and Biosynth Ltd for β-D-glucose-1-phosphate (MG02984).

### NMR

Dried extracts of parental and knockout bacterial cells were resuspended in 200 µL of a mixture of 90% D□O (Sigma-Aldrich) and 10% phosphate buffer (pH 7.4) prepared for NMR analysis as reported in Dona et al. (19). The solution was vortexed for one minute before centrifugation at 17,000 x g for 10 minutes at 4°C. An aliquot of 180 µL of the supernatant was transferred into a 3 mm NMR tube for NMR analysis.

All NMR spectra were acquired using a Bruker Avance III console combined with a 14.1 T magnet (600 MHz for ^1^H NMR). The system was equipped with a 5 mm broadband inverse (BBI) probe with z-axis magnetic field-gradient and a Bruker Sample Jet system, set to 3 mm shuttle mode with a cooling rack maintaining refrigerated tubes at 6° C.

A pre-saturated proton 1D-NOESY pulse sequence was used to obtain 1D ^1^H NMR spectra, with the offset set at 4.70 ppm for water suppression. 65,536 data points were collected with a spectral width of 20.0 ppm and a total of 64 scans (19).

A set of 2D NMR experiments were conducted for structural elucidation of the unknown compound present in the knockout samples, including Jres (J-resolved spectroscopy), ^1^H-^1^H-COSY (Correlation Spectroscopy) and ^1^H-^1^H-TOCSY (Total Correlation Spectroscopy). All NMR data were processed using TopSpin 3.6 (Bruker BioSpin GmbH, Rheinstetten, Germany). Processing steps included Fourier transformation, phasing, baseline correction, and spectral calibration.

Spiking experiments using an authentic chemical standard were used to further confirm that the unknown compound in the knockout sample was β-D-Glucose-1-Phosphate. The NMR sample of the knockout cells was spiked with increasing volumes of β-D-Glucose-1-Phosphate chemical standard (Biosynth Ltd; MG02984) ranging from 0.5 µL to 2 µL prepared at concentration of 30 µM in D_2_O, resulting in an increase in the signal intensity of the NMR resonances of interest in the knockout sample.

## Statistical analysis

The data are presented as the means±SDs from at least two biological replicates and two technical replicates per condition. Unpaired two-tailed Student’s *t*-tests were used to compare the data, and *p* < 0.05 was considered significant.

## Data availability

All data presented in this study are contained within the article and available from authors upon request to Gerald Larrouy-Maumus

## Supporting information

This article contains supporting information

## Biological safety considerations

The bacteria were handled within a Class-I safety-level cabinet.

## Acknowledgements

We would like to thank Dr Sufyan Pandor and Dr Hannah Florence from Agilent technologies for the discussion and advice on the separation and detection of sugar monophosphate by LC/Q-TOF.

## Author contributions

Yi Liu: investigation, data analysis, visualization, and writing–original draft preparation; Nadine Ruecker: investigation, data analysis;

Valwynne Faulker: investigation, data analysis, visualization; Kavitha Rachineni: investigation, data analysis, visualization; Nada Al-Saffar: investigation, data analysis, visualization; Pablo Villacampa: data analysis and editing;

Tiago Dias da Costa: supervision and writing-review and editing; Eachan Johnson: supervision and writing-review and editing;

Brian D. Robertson: investigation, funding acquisition, writing-review and editing; Sabine Ehrt: investigation, supervision, writing-review and editing;

Gerald Larrouy-Maumus: conceptualization, funding acquisition, data analysis, supervision, writing–original draft preparation and writing–review and editing.

All authors. All authors have read and approved the manuscript.

