## Supporting text for "Rv3400 is a phosphoglucomutase required for trehalose metabolism in *Mycobacterium tuberculosis*"

A comprehensive comparison of one-dimensional (1D) and two-dimensional (2D) NMR spectral data for both wildtype and knockout samples helped in identifying the unknown molecule present in the knockout sample.

Figure **S1** compares the 1D ^1^H-NOESY NMR spectra of the wildtype (blue) and knockout (red) samples, with spectral differences highlighted by red dotted rectangles. The knockout sample spectrum exhibits a distinct chemical shift resonance at 4.9 ppm, along with several additional resonances between 3.2 and 4.0 ppm, compared to the wildtype spectrum. To explore the correlations between these chemical shifts, 2D NMR spectra such as TOCSY and COSY were recorded. The observed complete end-to-end TOCSY correlations and COSY connectivity patterns indicate that these additional peaks in the knockout sample belong to a single molecule, suggesting that the unknown molecule may be sugar-related molecule. Notably, chemical shifts in the range of 5.5 to 3.0 ppm are commonly associated with sugar molecules.

Analysis of scalar coupling multiplets is essential to confirm the structure of this unknown sugar molecule. Therefore, 2D JResolved (JRES) spectra were also acquired. Figure **S2** compares the JRES spectra of the knockout and wildtype samples, with 1D spectra overlaid. Additional resonances in the knockout sample are highlighted with red circles. The scalar coupling multiplet pattern at 4.9 ppm, initially thought to be a triplet in the 1D NMR spectrum, is unexpectedly resolved into two doublets in the JRES spectrum. This splitting pattern resembles that observed for the anomeric protons of sugar phosphates such as glucose-1-phosphate, mannose-1-phosphate, and galactose-1-phosphate. Furthermore, as previously reported ([1](doi:%2010.1039/d0gc03290e%20(ref%20for%20glucose-1,6-bisphosphate))), a six-membered sugar exhibiting a triplet peak at 4.91 ppm has been identified as glucose-1,6-bisphosphate. A comparable doublet pattern was observed at 4.0 ppm for protons on 6^th^ carbon of glucose-6-phosphate in the JRES spectrum which is attributed to ^1^H-^31^P scalar coupling ([2](https://chemistry-europe.onlinelibrary.wiley.com/doi/10.1002/cphc.202200495)). A similar doublet pattern observed in the knockout sample's JRES spectrum at a chemical shift position 4.9 ppm suggests the presence of **¹**H–¹H coupling along the indirect dimension and ¹H–³¹P coupling along the direct dimension, indicating that the molecule is a sugar with a phosphate group.

The presence of six additional resonances suggests that the unknown compound may be a six-membered sugar phosphate (pyranose form). Further analysis of the scalar coupling constants, ranging from 8.0 to 15.0 Hz (as observed in the JRES spectrum), indicates that all protons in this unknown compound are axially oriented relative to each other ([3](https://pubs.rsc.org/en/content/articlelanding/1971/j2/j29710000446/unauth)). This observation supports the hypothesis that the sugar moiety of the unknown molecule should be glucose, as glucose typically exhibits an axial orientation of all its protons. The overall chemical shift assignment of the unknown molecule in the knockout sample suggests that the resonance at chemical shift 4.9 ppm corresponds to the anomeric proton of a glucose molecule. The JRES spectrum further supports the presence of a phosphate group on the first carbon (the anomeric carbon), indicating that the unknown molecule is an anomer of glucose-1-phosphate. However, glucose-1-phosphate can exist in either α- or β- anomeric form. In this case, the anomeric proton exhibits a vicinal coupling constant of 7.80 Hz, which is characteristic of the β-anomer of glucose-1-phosphate, confirming the identity of the unknown molecule in the knockout sample.

Spiking studies were performed using a standard solution (36 mM) of β-D-glucose-1-phosphate. In each spiking point, 0.5 μL of the standard solution was added to the knockout sample. The spiking experiment confirmed that the unknown molecule in the knockout sample is β-glucose-1-phosphate, as clearly demonstrated in **S3** with the required spectral expansions. For further verification, 2D JRES, COSY, and TOCSY NMR spectra of authentic β-D-glucose-1-phosphate chemical standard were compared with those of the knockout sample (Figures **S4** and **S5.**). This comparison revealed identical scalar coupling multiplet patterns and chemical shift correlations between the two samples, unambiguously confirming that the unknown molecule in the knockout sample is β-glucose-1-phosphate.

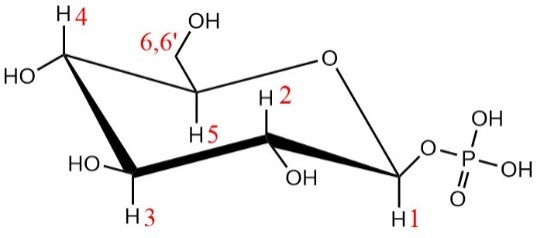

β-D-glucose-1-phosphate

**S.Table :1.** Chemical shift values of unknown molecule that match those of β-D-Glucose-1-Phosphate

| **Peaks** | **^1^H Chemical shift δppm** | **Number of Hs** | **Spin Type** | **Multiplicity** | | **Correlations** | |
| --- | --- | --- | --- | --- | --- | --- | --- |
|  |  |  |  | **Type** | **Coupling (*^3^J_HH/_* Hz)** | **COSY (ppm)** | **TOCSY (ppm)** |
| 1 | 4.91 | 1 | CH | 2d | *^3^J_H1-H2_* _=_7.80  *^4^J_H1-P_* _=_7.80 | 3.32 | 3.32, 3.35, 3.54 |
| 2 | 3.32 | 1 | CH | dd | *^3^J_H1-H2_* _=_7.80,  *^3^J_H2-H3_* _=_9.30 | 3.54, 4.91 | 4.91, 3.35, 3.54, 3.51, 3.68, 3.92 |
| 3 | 3.54 | 1 | CH | t | *^3^J_H2-H3_* = 9.30,  *^3^J_H3-H4_* _=_9.68 | 3.32, 3.35 | 4.91, 3.32, 3.35, 3.51, 3.68, 3.92 |
| 4 | 3.35 | 1 | CH | t | *^3^J_H3-_*_H4_ & *^3^J_H4-_*_H5 =_ 9.70 | 3.51, 3.54 | 4.91, 3.32, 3.35, 3.54, 3.68, 3.92 |
| 5 | 3.51 | 1 | CH | dd, dd | *^3^J_H5-H6_* _=_ 6.91,  *^3^J_H4-H5_* _=_9.70,  *^3^J_H5-H6’_* _=_ 2.25 | 3.35, 3.68 | 3.68, 3.54, 3.35, 3.32 |
| 6 | 3.68 | 2 | CH2 | dd | *^3^J_H5-H6_* _=_ 6.91,  *^3^J_H6’-H6_* _=_ 12.30 | 3.51, 3.92 | 3.92, 3.54 3.51, 3.35, 3.32 |
| 6’ | 3.92 | 2 | CH2 | dd | *^3^J_H5-H6’_* _=_ 2.25,  *^3^J_H6-H6’_* _=_ 12.30 | 3.51,3.68 | 3.68, 3.54, 3.35, 3.51, 3.32 |

*doublet of doublet (dd); triplet (t); doublet(d)*

References

1. DOI: 10.1039/d0gc03290e

2. <https://doi.org/10.1002/cphc.202200495>
3. <https://pubs.rsc.org/en/content/articlepdf/1971/j2/j29710000446>
