## Supporting Figures for "Rv3400 is a phosphoglucomutase required for trehalose metabolism in *Mycobacterium tuberculosis*"

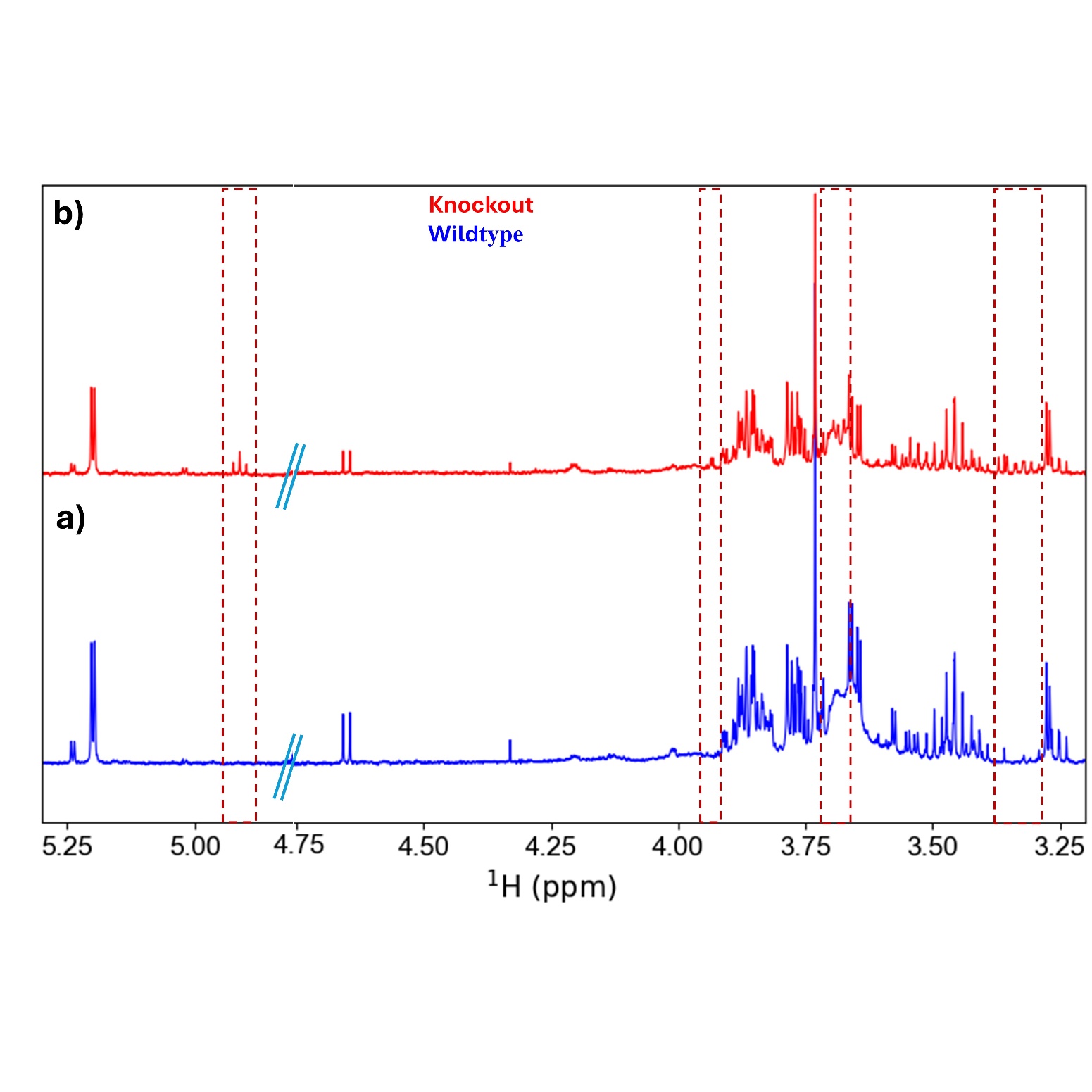

**S1:** 1D ^1^H NOESY spectral comparison of (a) Wildtype and (b) Knockout samples. Rectangular dotted boxes indicate additional resonances observed in the Knockout samplethat are absent in the Wildtype sample. The crossed blue line at 4.7 ppm marks removed water signal.

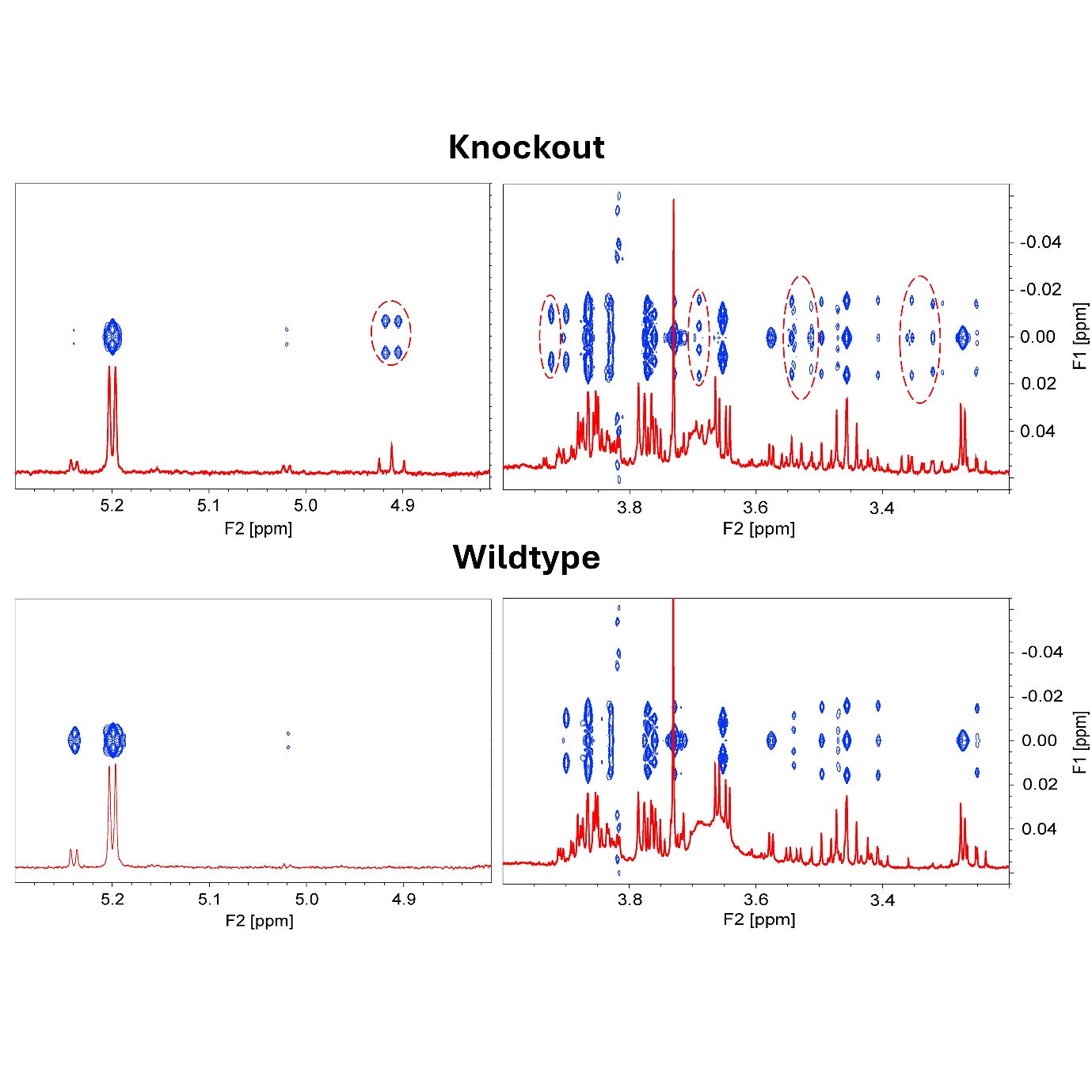

**S2:** 2D JRES spectra (blue) overlaid with 1D ^1^H NMR spectra (red ) from both Wildtype and Knockout samples. Red circles highlight the presence of additional spectral resonances present in the Knockout samples but absent in the Wildtype.

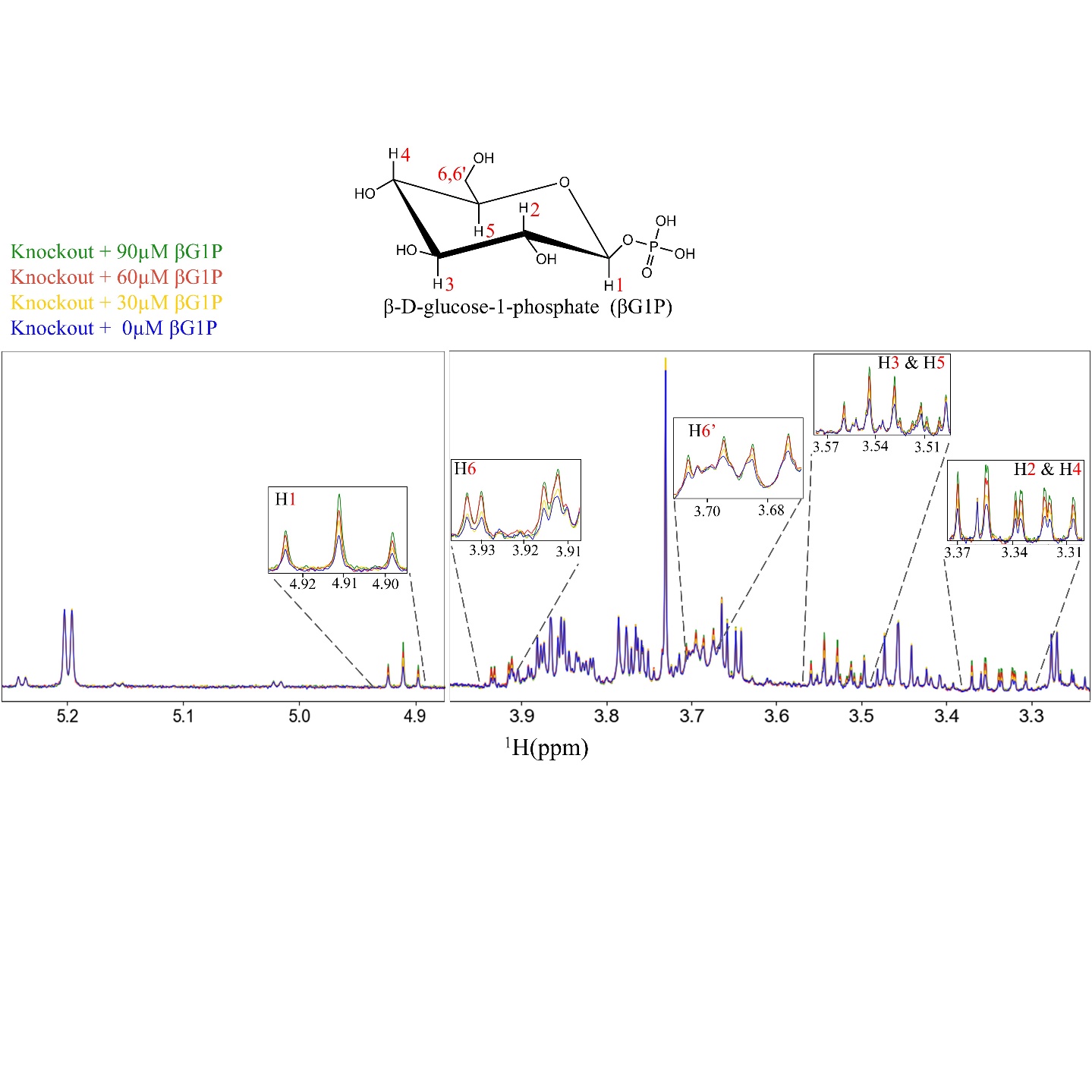
**S3:**  1D ^1^H NOESY NMR spectral overlay showing the spiking experiment results. The blue spectrum represents the Knockout sample without spiking, while the yellow, red, and green spectra correspond to the samples spiked with β-D-glucose-1-phosphate at concentrations of 30 µM, 60 µM, and 90 µM, respectively. Assignments for all the protons of β-D-glucose-1-phosphate are provided with spectral expanisons.

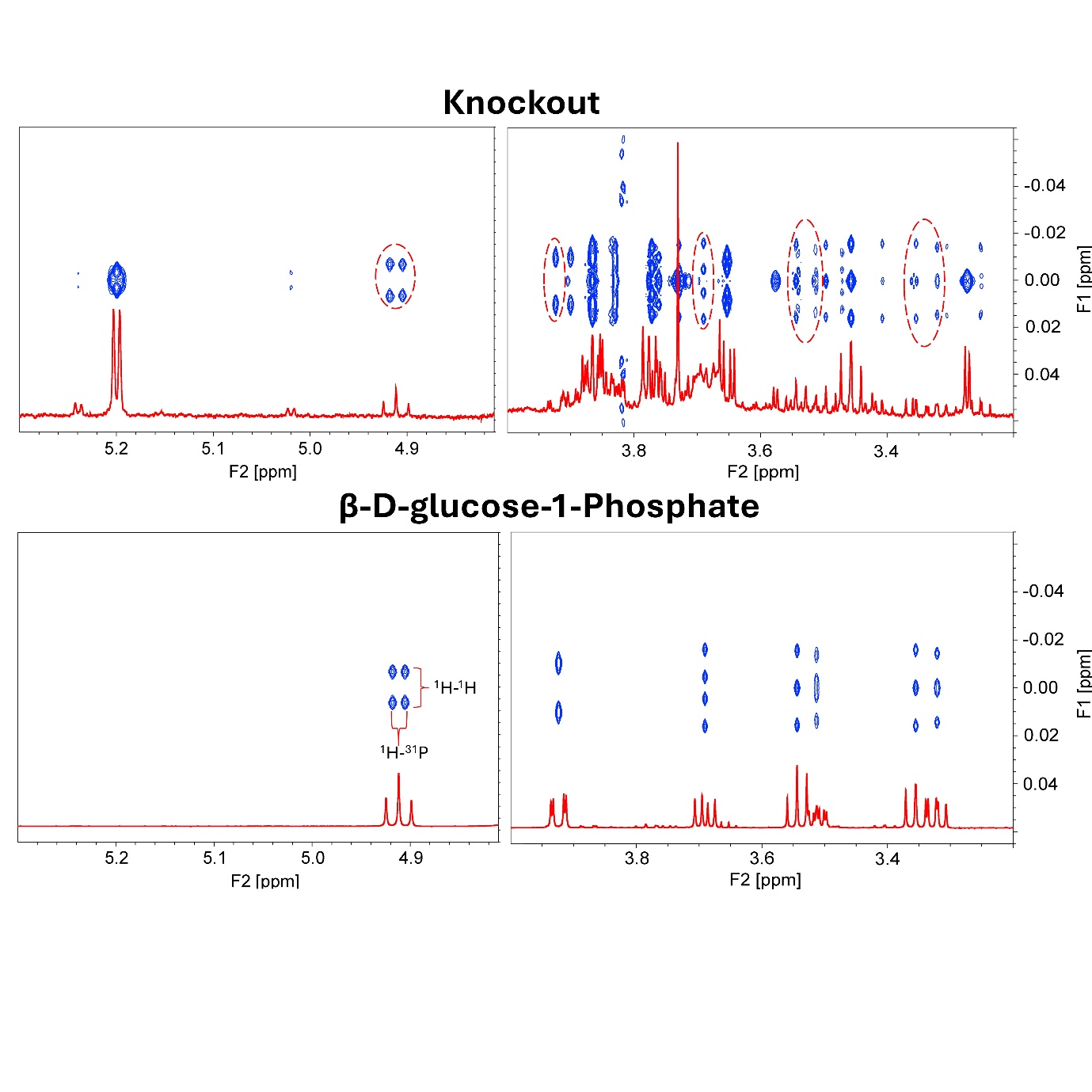

**S4:** 2D JRES spectra (blue) overlaid with 1D ^1^H NMR spectra (red) from the Knockout sample and the chemical standard β-D-Glucose-1-Phosphate. Red circles in the Knockout sample spectra highlight resonances that are identical resonances to those observed in the chemical standard β-D-glucose-1-phosphate spectra.

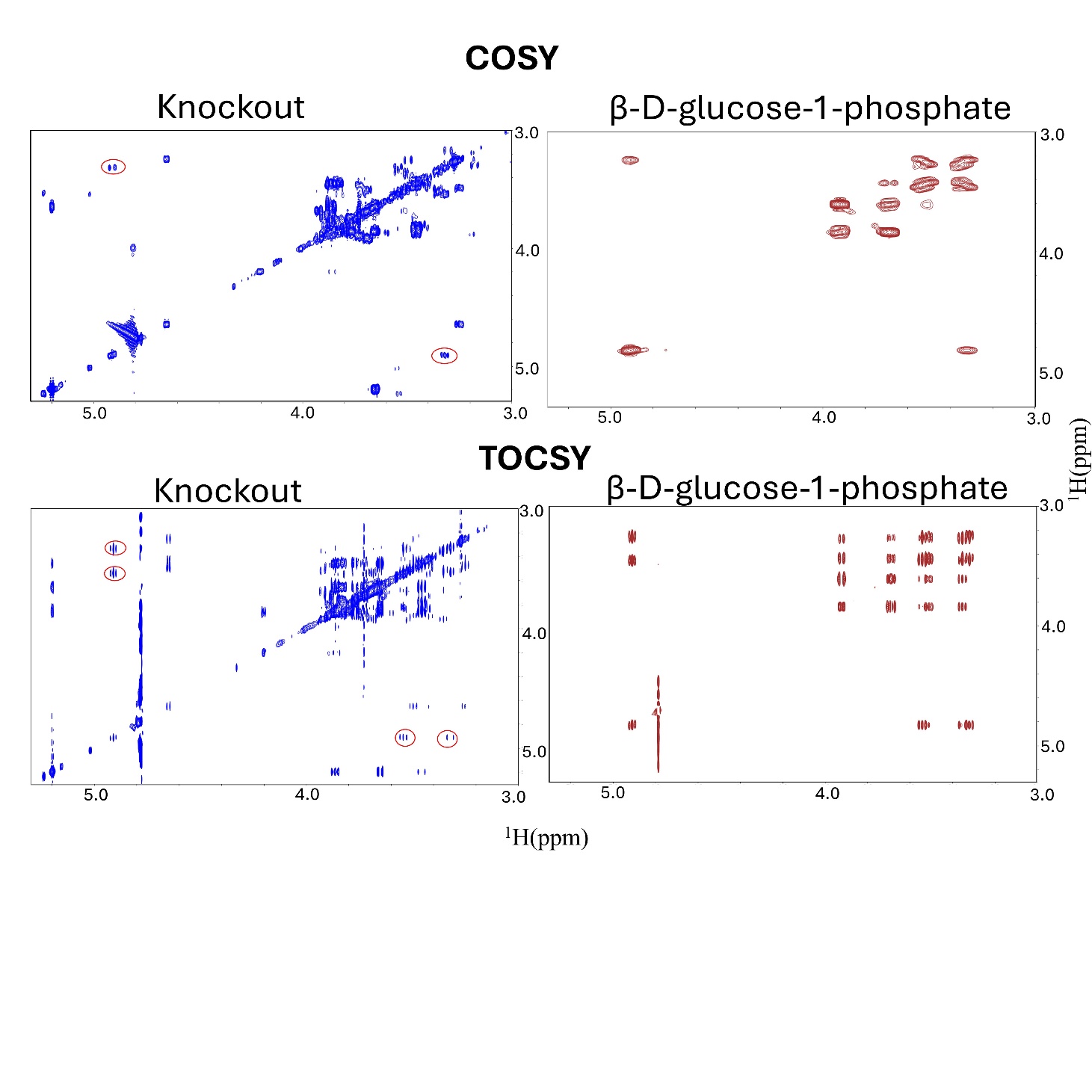
**S5:** Comparison of 2D COSY and 2D TOCSY NMR spectra between the Knockout sample and β-glucose-1-Phosphate. The figures highlight the presences of β-D-Glucose-1-Phosphate resonances in the Knockout sample, indicated by red circles

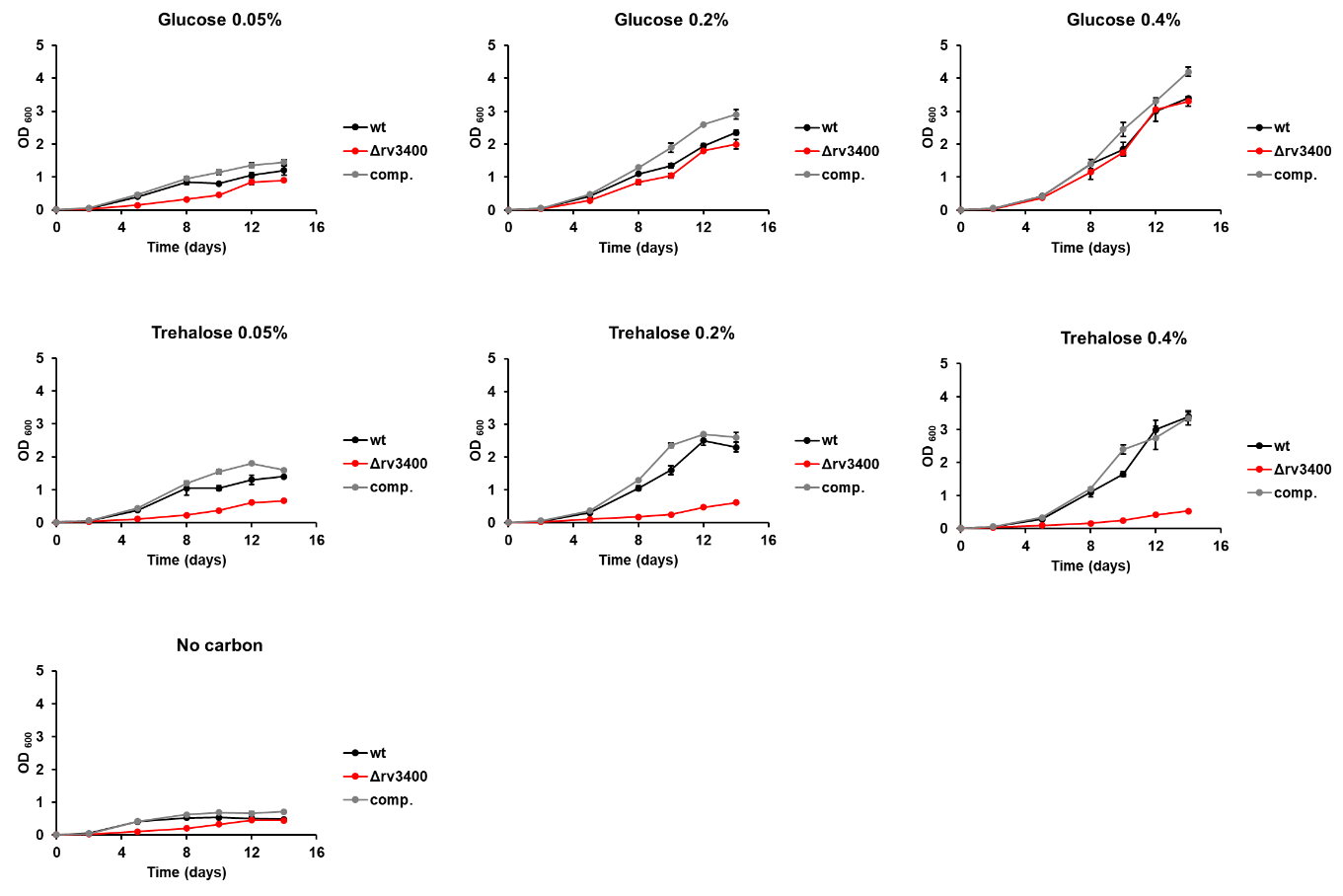

**S6:** Growth curves of parental (black), Δrv3400 (red), complemented (grey) strains with no carbon source, 0.05%, 0.2% or 0.4% glucose or trehalose as the sole carbon source and using standard BSA.

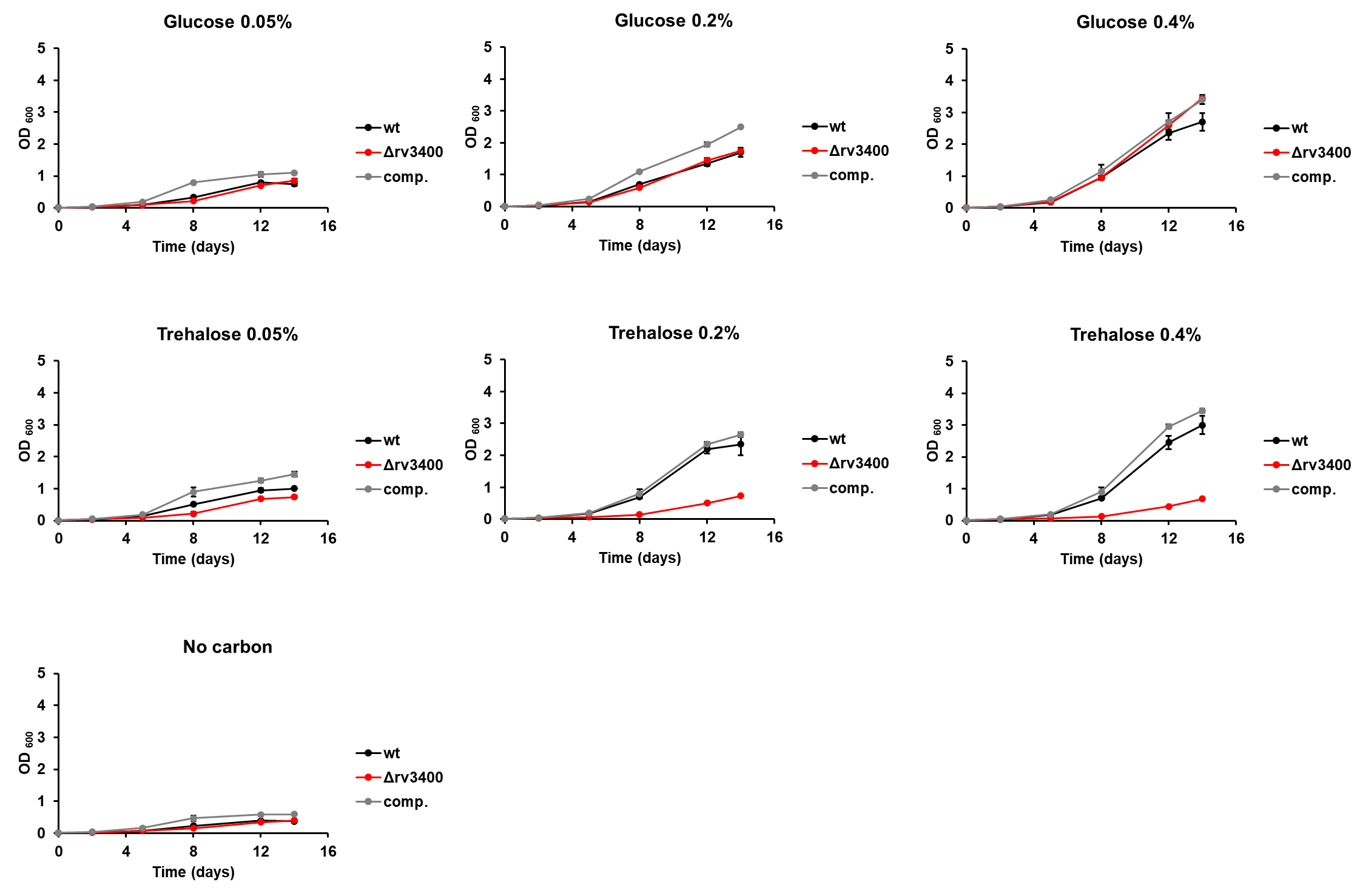

**S7:** Growth curves of parental (black), Δrv3400 (red), complemented (grey) strains with no carbon source, 0.05%, 0.2% or 0.4% glucose or trehalose as the sole carbon source and using BSA fraction V fatty acid free.
